# Single cell transcriptome mapping identifies a local innate B cell population driving local antibody production and chronic rejection after lung transplantation

**DOI:** 10.1101/2021.02.16.431278

**Authors:** Natalia F. Smirnova, Kent Riemondy, Susan Collins, Kapil N. Patel, Carlyne Cool, Melanie Koenigshoff, Nirmal S. Sharma, Oliver Eickelberg

## Abstract

Bronchiolitis obliterans syndrome (BOS) due to chronic rejection is the main reason for poor outcomes after lung transplantation (LTx). We and others have recently identified B cells as major contributors to BOS after LTx. The extent of B cell heterogeneity, however, as well as those B cell populations that determine chronic rejection of the lung, remain entirely unclear. Here, we provide a map of cell population heterogeneity and their gene expression patterns during chronic rejection after orthotopic LTx in mice. Out of a total of 14 major comprehensive cell clusters, corresponding to 11 known major cell subpopulations, *Mzb1*-expressing plasma cells (PCs) were the most prominently increased cell population in lungs with BOS. *Scgb1a1*-expressing bronchial epithelial cells were depleted, while *Cd14*-expressing monocytes and *Pdgfra*-expressing fibroblasts were enriched in BOS grafts. These findings were validated in two different cohorts of human BOS after LTx. MZB1-IgG-double positive PCs infiltrated the airways of patients with BOS, while they were absent from healthy lungs. IgG, but not IgA, IgD, IgE or IgM, were significantly increased in bronchoalveolar lavage from BOS patients, compared with LTx patients who did not develop BOS. Pseudotime and trajectory analysis revealed that a *Bhlhe41*, *Cxcr3, Itgb1*-triple positive-B cell subset, also expressing classical markers of the innate-like B-1 B cell population, served as the progenitor pool for *Mzb1*+ PCs. This progenitor B cell subset accounted for the increase in IgG_2c_ expression and production in BOS lung grafts. Importantly, *Aicda^−/−^* mice, which lack all isotypes of Ig (except IgM) were protected from the onset of BOS after LTx. In summary, we provide a detailed atlas of cell population changes of chronic rejection after LTx. We have identified IgG-positive PCs and their progenitors – an innate B cell subpopulation characterized by a specific subset of markers - as the major source of local antibody production and a major contributor to both mouse and human BOS after LTx.

## Introduction

Chronic lung diseases (CLD) are a leading cause of morbidity and mortality worldwide, accounting for 7% of all deaths from noncommunicable diseases (1). To date, lung transplantation (LTx) represents the only potential cure for most end stage CLD. While short-term survival after LTx has significantly improved over time, the median and long-term survival of LTx is severely limited primarily due to chronic rejection of the allograft, which leads to an irreversible decline of lung function. Adults who underwent primary LTx in the recent era (2010 - June 2017) exhibited a median survival of 6.7 years, the worst of any solid organ transplantation (2). The long-term outcome of LTx is mainly limited by the development of Bronchiolitis Obliterans Syndrome (BOS), the most common phenotype of chronic rejection. BOS develops in 50% of all patients 5 years after LTx, and is the leading cause of mortality after receiving a lung transplant (2). During the development of BOS, airways are progressively infiltrated by leukocytes, exhibiting signs of chronic inflammatory and immune processes, ultimately resulting in peribronchial fibrosis and the loss of the bronchial epithelium. This scarring, due to epithelial loss, leads to an obstruction of the airways, decreasing the diameter of their lumen, limiting airflow, and reducing lung function (3).

We and others recently reported that B cells are a major contributing cell type to the development of BOS (4, 5), which has been reported in chronic rejection of other solid organs after transplantation (6). Most of these studies have characterized systemic or local B cells in chronic rejection using flow cytometry (7) or immunostaining of tissue sections (4, 5), introducing an intrinsic bias, due the use of prespecified surface markers. The advent and validation of new methods, such as single cell RNA-sequencing (scRNA-seq), offers the possibility to avoid this bias, by distinguishing cell populations based on an entire set of transcripts expressed by each cell. The scRNA-seq technology has already changed preexisting classifications of immune cell subsets (8), and provided a new level of understanding of their distribution and function in disease (9, 10).

Here, we sought to provide an atlas of BOS after LTx using scRNA-seq to identify changes in the composition of cell types present during rejection inside the lung grafts, and alternations to their individual gene expression profiles. Plasma cells (PCs) were the most enriched cell cluster detected in BOS. PCs were consistently found in large amounts in subepithelial layers of human and mouse BOS airways. Detailed subclassification analysis of B cells and their progeny in murine lung grafts identified a subset of B cells that give rise to PCs. These B cells expressed a number of markers of the innate B-1 cell population and were capable of producing IgG *ex vivo*. Mechanistic interrogation of these findings *in vivo* revealed that a mouse strain lacking production of all but IgM immunoglobulin isotypes (*Aicda^−/−^* mice) displayed protection against the development of BOS after LTx.

## Results

### Single cell RNA-sequencing identifies BOS-associated cell populations

B6 recipient mice underwent orthotopic left lung transplantation (LTx) lungs from B6 (control) of HLA (chronic rejection) donors (syngeneic group: B6➝B6, *n*=5; and BOS group: HLA➝B6, *n*=3), as previously described (4), to generate robust lymphocytic bronchiolitis and BOS. Lung grafts harvested at 1 month after LTx (**Fig. 1A**) were dissociated using a combination of enzymatic and mechanical methods, after which total cells were sorted. After application of quality control filters and removal of ambient RNA, 9,697 cells from syngeneic grafts and 11,081 cells from BOS grafts were included in scRNA-seq analysis (**Sup. Fig. 1A**: detailed composition of cells per sample). Cellular gene expression data from both conditions were aligned and projected in a bidimensional UMAP plot to identify cell populations present in the lung grafts. Unsupervised clustering, using the Seurat package, distinguished 14 distinct cell clusters, based on their individual gene expression profiles (**Fig. 1B**). Putative biological identities were assigned to each cluster using patterns of established canonical markers, as well as importing the murine gene expression atlas, which resulted in the identification of 11 cell types (Tabula Muris (11), **Sup. Fig. 1**). As shown in **Fig. 1B**, lung grafts were composed of structural resident, as well as infiltrating or resident immune and inflammatory cells. Major shifts in cell abundance of each cluster were observed between the syngeneic and BOS groups (**Fig. 1C**). The most prominently increased cell type in chronic rejection were plasma cells (PCs), defined by the expression of *Xbp1*, *Prdm1* (encoding BLIMP-1), *Sdc1* (encoding CD138), and *Mzb1*(12, 13) (**Fig. 1C**). Based on this finding, and on our previous data demonstrating a major role of B cells in the development of BOS(4), we next sought to define this cluster in more detail and assess the mechanistic contribution and clinical relevance of B cell subpopulations and plasma cells in BOS.

**Figure 1.**
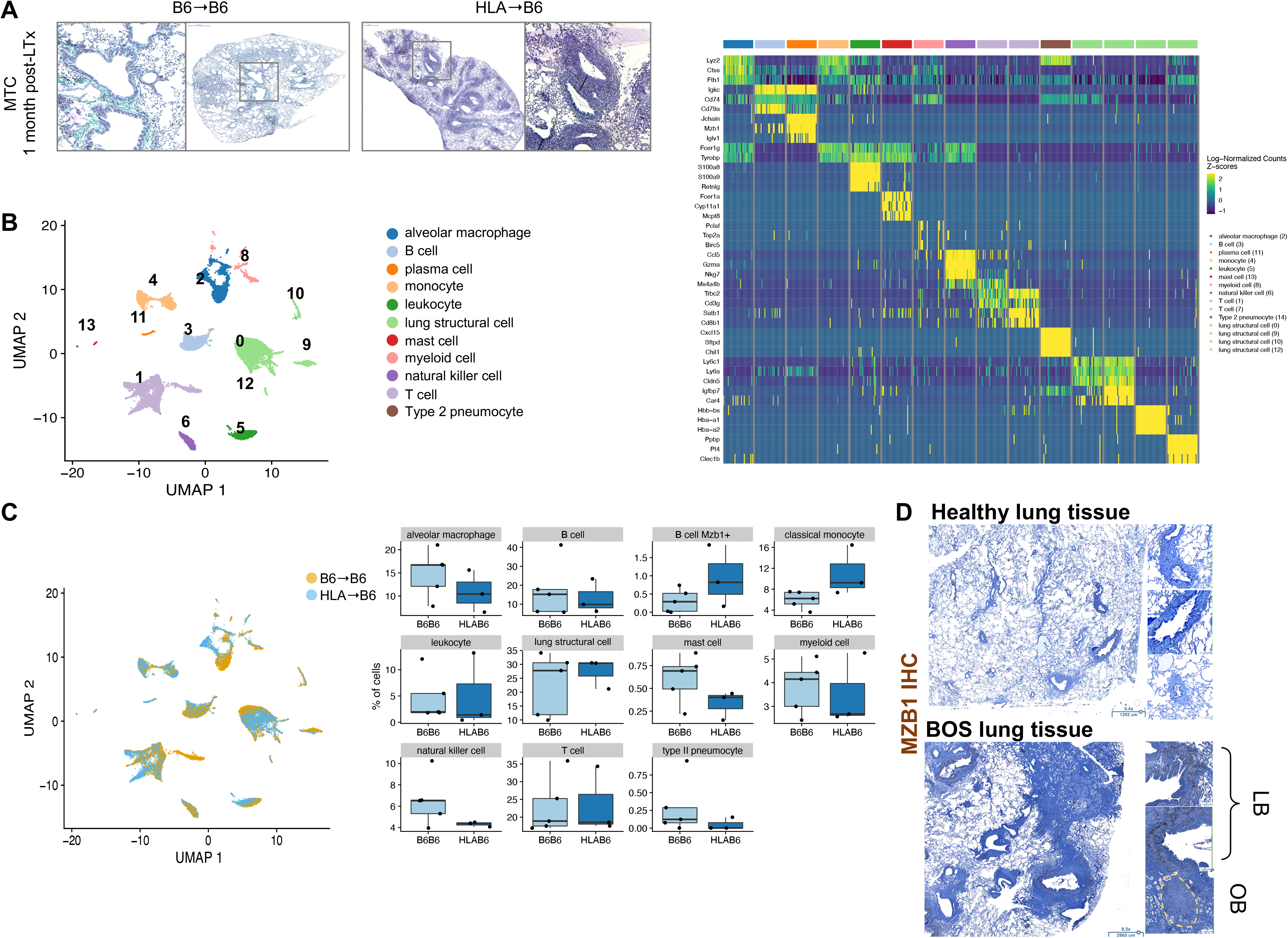
Single cell RNA-sequencing identifies BOS-associated cell populations. **A,** Representative histology sections of control (B6➝B6) and BOS (HLA➝B6) mouse lung grafts one month after lung transplantation (LTx). Staining: Masson Trichrome. **B,** Uniform Manifold Approximation and Projection (UMAP) representation of 11 distinct cell populations from unsupervised clustering of gene expression data in single cells extracted from control (B6➝B6) and BOS (HLA➝B6) transplanted mouse lungs (**left**) and heat map showing most upregulated genes for each of the cell clusters (**right**)**. C,** UMAP representation of the cell populations detected in murine lung grafts by condition (HLA➝B6 and B6➝B6). Proportion of each cell type in control vs. BOS mouse lung graft samples (**right**). **D,** Immunohistochemistry (IHC) staining for MZB1 on explants from healthy donors and patients who developed BOS after LTx (samples from the University of Colorado collection). LB: lymphocytic bronchiolitis, OB: obliterative bronchiolitis.

Of note, the resident structural cell compartment was subdivided into 11 distinct clusters, which we annotated manually (**Sup. Fig. 2**). Major quantitative shifts were observed for structural clusters #0 to #3. Cluster #0, enriched in BOS grafts, was identified as lymphatic endothelial cells, based on their positivity for *Tmem100* and *Lyve1*(14). Interestingly, and consistently with its association with rejection, endothelial cells in this cluster were highly positive for genes encoding MHC II molecules. Expression of MHC II is enhanced on endothelial cells in inflammatory contexts, conferring them an antigen presenting role (15). Consistently with the pathological changes documented for BOS (16, 17), cluster #1 and #2 cells, expressing lung epithelial cell markers *Scgb1a1* and *Sftpc* were decreased in proportion in rejecting lungs(16, 18), while cluster #3 *Col1a1*-expressing fibroblasts were enriched (19)(**Sup. Fig. 2**).

**Figure 2.**
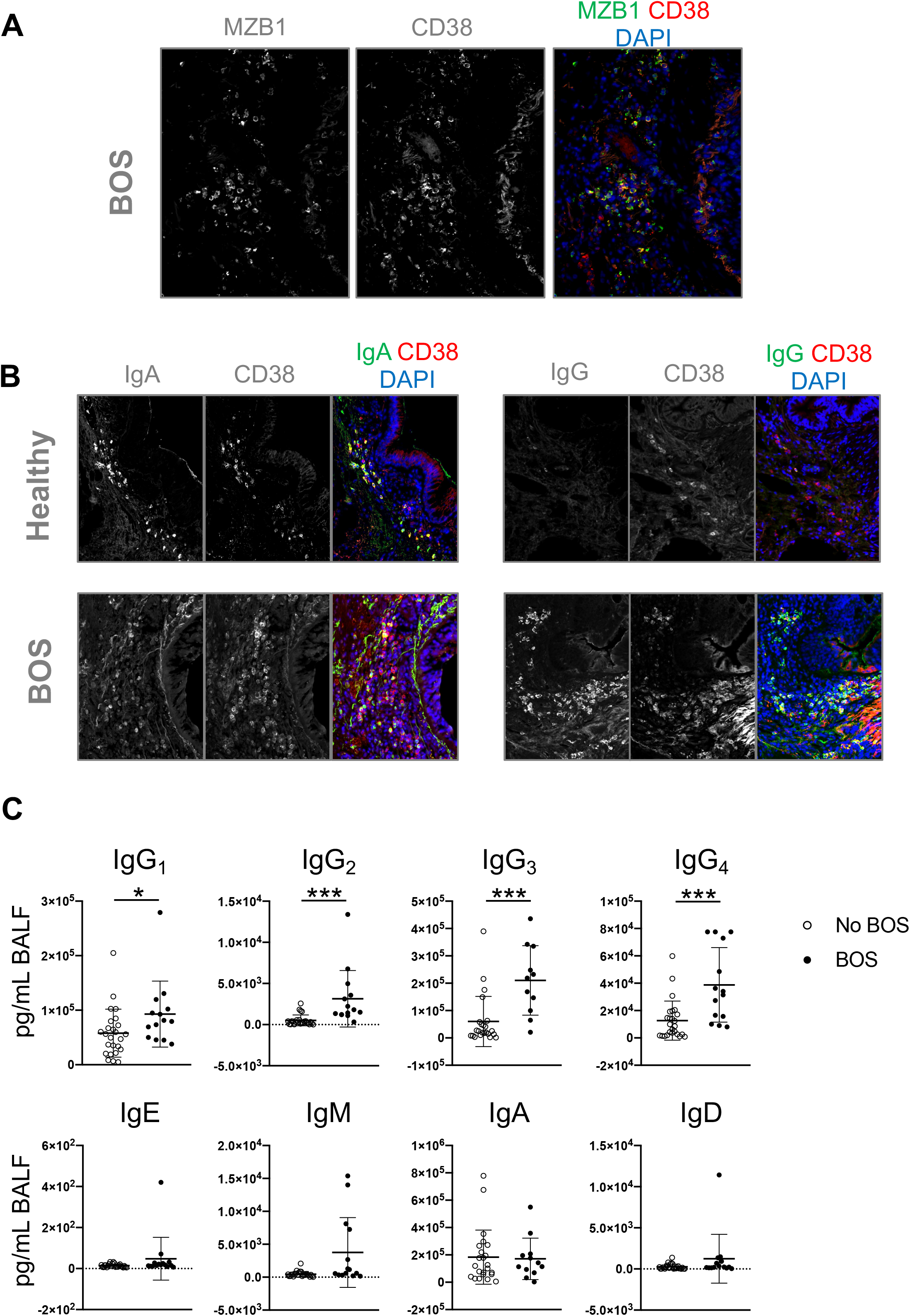
IgG-positive PCs and increase in local IgG are signatures of human BOS. **A,** Immunofluorescence (IF) staining of MZB1 and CD38 on a lung tissue sample with BOS after LTx (samples from the University of Colorado collection). **B,** IF staining of IgA and CD38 in healthy donor and BOS tissue samples (**left**) and staining of IgG and CD38 in healthy and BOS tissue samples (**right**) (samples from the University of Colorado collection). **C,** Quantification of the local Ig profile in BALF samples from transplanted patients with and without BOS after LTx (USF-TGH LTx cohort). Data are presented as mean+/−SEM, and analyzed with a Mann-Whitney unpaired t-test. \**p*<0.05; \*\*\**p*<0.001.

### Plasma cells and immunoglobulin production in human lung grafts with BOS

To assess the clinical relevance of the increased PC population in BOS, we compared explants from patients, who developed BOS grade 3 and underwent re-transplantation, with healthy controls (n=10 each). PCs, identified by the expression of MZB1, were robustly detected in large amounts in the peribronchial areas in BOS (**Fig. 1D**). While these peribronchial PCs displayed similar immunoglobulin A (IgA) expression in healthy and BOS samples, numerous IgG-positive PCs were observed around the airways in BOS, while undetectable in healthy samples (**Fig. 2A, 2B**). Quantification of local Ig isotype production in the bronchoalveolar lavage fluid (BALF) samples from transplanted patients with or without BOS (**Table 1**) showed that the titers of all four IgG subclasses (IgG_1_, IgG_2_, IgG_3_, IgG_4_) were significantly increased in BOS, while no differences were detected for IgE, IgM, IgA, or IgD isotypes comparing LTx patients with and without BOS (**Fig. 2C**).

**Table 1:**
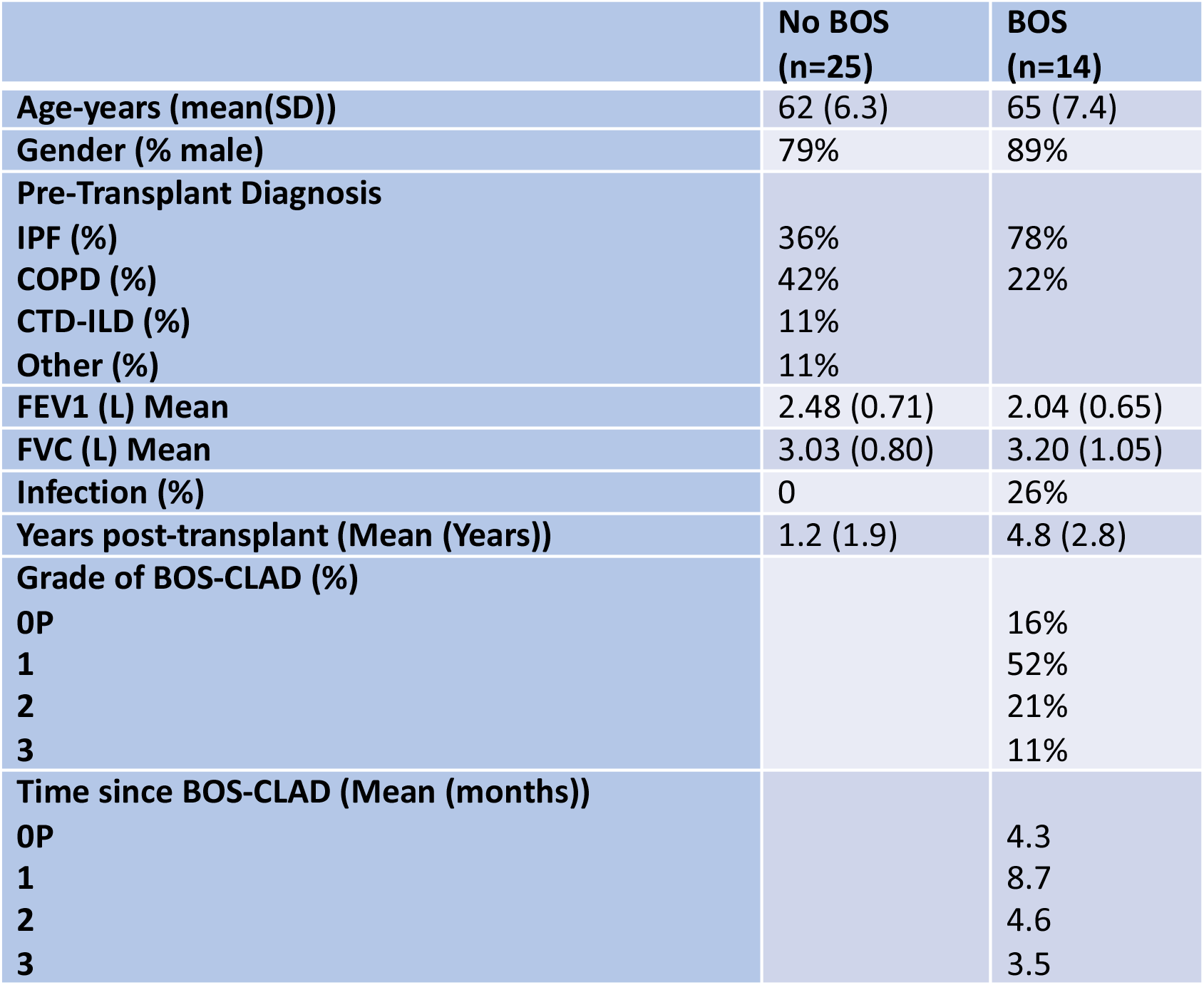
TGH/USF patients demographics

Importantly, RNA-sequencing of frozen lung tissue from explants with BOS (*n*=3) revealed that *Mzb1* and *Cxcl13*, encoding a chemokine primarily known for its involvement in B cell homing, were amongst the highest and most significantly upregulated genes in BOS compared to control lung samples (increased by 36.50-and 47.10-fold, respectively; \**p*<0.05; *data not shown*).

### Heterogeneity of B cells in lung grafts

An unsupervised detailed analysis of B and PC populations in mouse lung grafts identified four distinct subpopulations: three B cell clusters (clusters #0, #1, and #3) and a PC cluster (cluster #2) (**Fig. 3A, 3B**). The majority of cells were found in cluster #0, and their amounts were similar between controls and BOS (226 and 254 cells, respectively, in B6➝B6 and HLA➝B6 lung grafts). Cell numbers in clusters #1 and #3 were increased 4-and 3-fold in HLA➝B6 versus B6➝B6, respectively. Cluster #2, identified as the PC cluster, based on the expression of *Xbp1*, *Mzb1*, *Sdc1*, *Prdm1*, displayed a 10-fold-increase in cell numbers in HLA➝B6 versus B6➝B6 lung grafts (**Fig. 3C**). Pseudotime and velocity analysis were used to interrogate the origin of plasma cells in the HLA➝B6 lungs showing BOS (**Fig. 3D**). Pseudotime analysis, using CytoTrace (20), which identifies stem and differentiated cells based on the total numbers of genes expressed, showed that PCs express fewer genes, while cluster #1 B cells expressed the most genes compared to the general population. This suggests that cluster #2 PCs are in a highly differentiated state, while cluster #1 cells are likely a progenitor population. The velocity analysis showed a directionality between cluster #1 and cluster #2 PCs, suggesting that PCs originate from cluster #1 B cells. The gene *Bhlhe41* was one of the predominant markers of cluster #1 cells. We validated by immunostaining the infiltration of BHLHE41-positive cells around the airways of both humans and mice who developed BOS after LTx (**Fig. 3E**).

**Figure 3.**
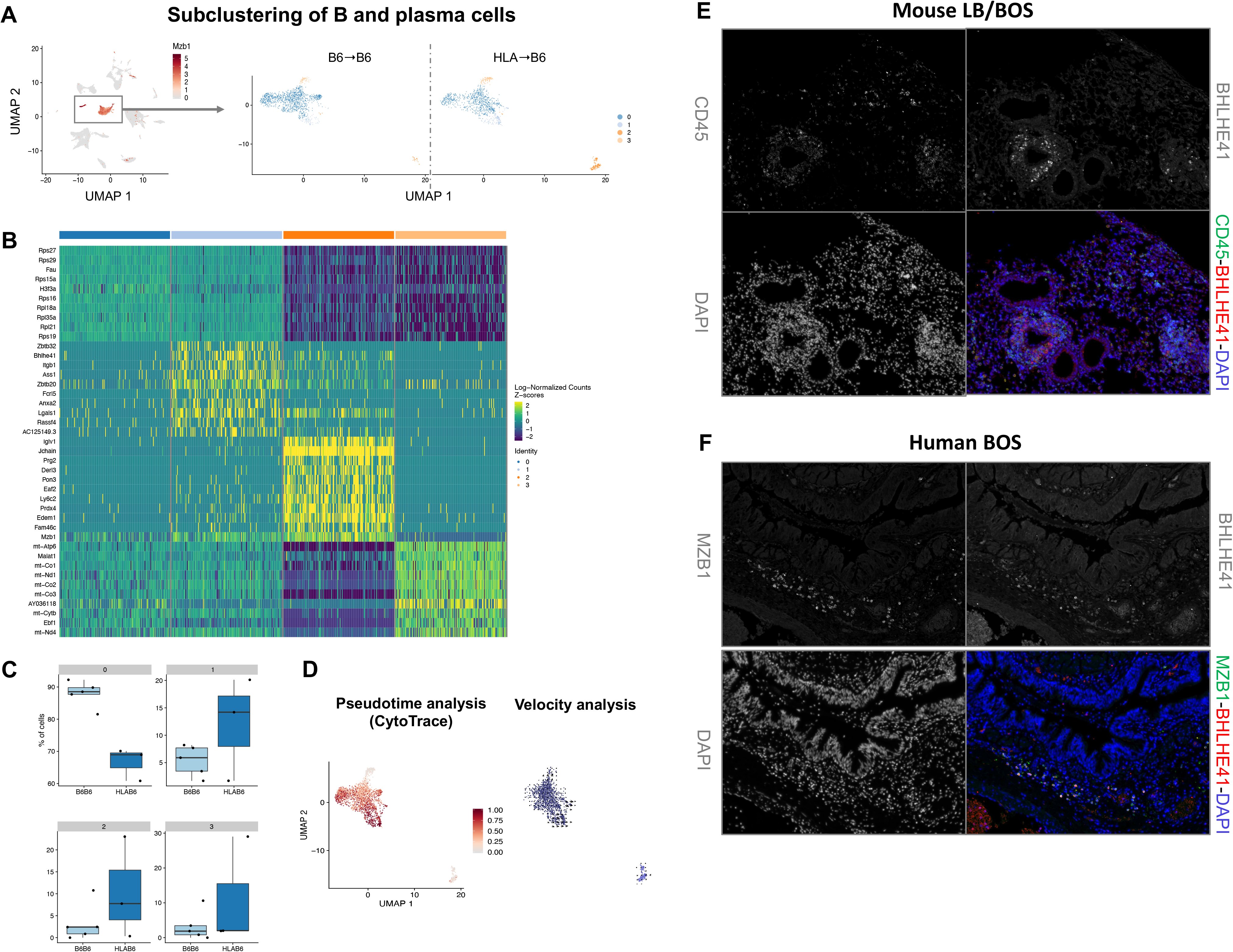
Heterogeneity of B cells and their progeny in the lung graft. **A,** Unsupervised clustering of B and plasma cells in control (B6➝B6) and BOS (HLA➝B6) mouse lung grafts, 1 month after lung transplantation (LTx). **B,** Heat map showing most upregulated genes in the B and plasma cell clusters. **C,** Proportion of each cell type in control (B6➝B6) and BOS (HLA➝B6) lung grafts. **D,** CytoTrace pseudotime analysis on the B and plasma cell subclusters comparing total number of genes expressed in each cluster (**left**) and Velocity analysis showing directionality from cluster 1 B cells to cluster 2 plasma cells (**right**). **E,** Immunofluorescence (IF) staining of the hematopoietic marker CD34 and BHLHE41 in lung graft tissue from a BOS (HLA➝B6) mouse (**top**), and MZB1 and BHLHE41 in a lung tissue sample from a patient with BOS after LTx (samples from the University of Colorado collection) (**bottom**).

### Functional characterization of the cluster #1 B cell subset

To better understand the origin and functions of PCs in BOS, we investigated the cells from cluster #1, identified as potential progenitors of PCs in the BOS lung graft. Cluster #1 B cells expressed a number of transcription factors: *Zbtb32*, described in memory B cells, but absent in PCs, *Zbtb20*, expressed in B-1 cells and peaking in PCs (21), and *Bhlhe41*, recently unraveled as a marker of the B-1 population, which regulates B-1 cell differentiation, homeostasis and function (22). Based on its first 50 marker genes, cluster #1 B cells were defined as an innate B-1 cell population, according to the ImmGen database (23). Cluster #1 was in addition characterized by the expression of the surface markers *Cxcr3*, and *Itgb1*, respectively involved in lymphocyte homing and adhesion, and without any previously reported functional role in B-1 cells (**Fig. 4A**). The expression of *Cxcr3* and *Itgb1* was significantly higher in cluster #1 compared to the others, with a respectively 6 and 7-fold higher expression in cluster #1. We used these surface markers, combined to the B cell marker CD19 in flow cytometry, to distinguish cluster #1 cells (CD19+CXCR3+ITGB1+) from the other clusters (CD19+CXCR3-ITGB1-). At 1 month after LTx, HLA➝B6 lung grafts presented an increase in total CD19+ B cells, as well as in CD19+IgG+ B cells (**Sup. Fig. 3**). To understand which cell population accounted for this increase in IgG+ cells, we measured surface IgG amongst the CD19+CXCR3+ITGB1+ versus CD19+CXCR3-ITGB1-populations and found that the CD19+CXCR3+ITGB1+ cells specifically displayed an increase in surface IgG in HLA➝B6 versus B6➝B6 lung grafts (**Sup. Fig. 3**). All Ig isotypes, with the exception of IgD, were expressed in cluster #1 and PCs on a gene level (**Fig. 4B**). The absence of expression of IgD in cluster #1 cells is consistent with a B-1 phenotype. To determine if cluster #1 B cells were capable of secreting Ig, we sorted these cells from mouse lungs with or without BOS 1 month after LTx, and put them in culture with LPS (1 ug/mL) stimulation. A phenotypic analysis by flow cytometry confirmed that cluster #1 cells express the B-1a cell markers CD43 and CD5 (**Sup. Fig. 3**). Cluster #1 cells secreted significantly higher amounts of IgG_2c_ unlike the other B cells, and this secretion was increased when the cells were sorted from lungs with BOS compared with controls (**Fig. 4C**). The secretion of IgG_2c_ was also significantly increased in the BALF collected from lung grafts from HLA➝B6 versus B6➝B6 mice (**Fig. 4D**). Altogether, we identified a B cell population with B-1 markers, which infiltrates rejected lung grafts and is accountable for an increased local production of IgG_2c_.

**Figure 4.**
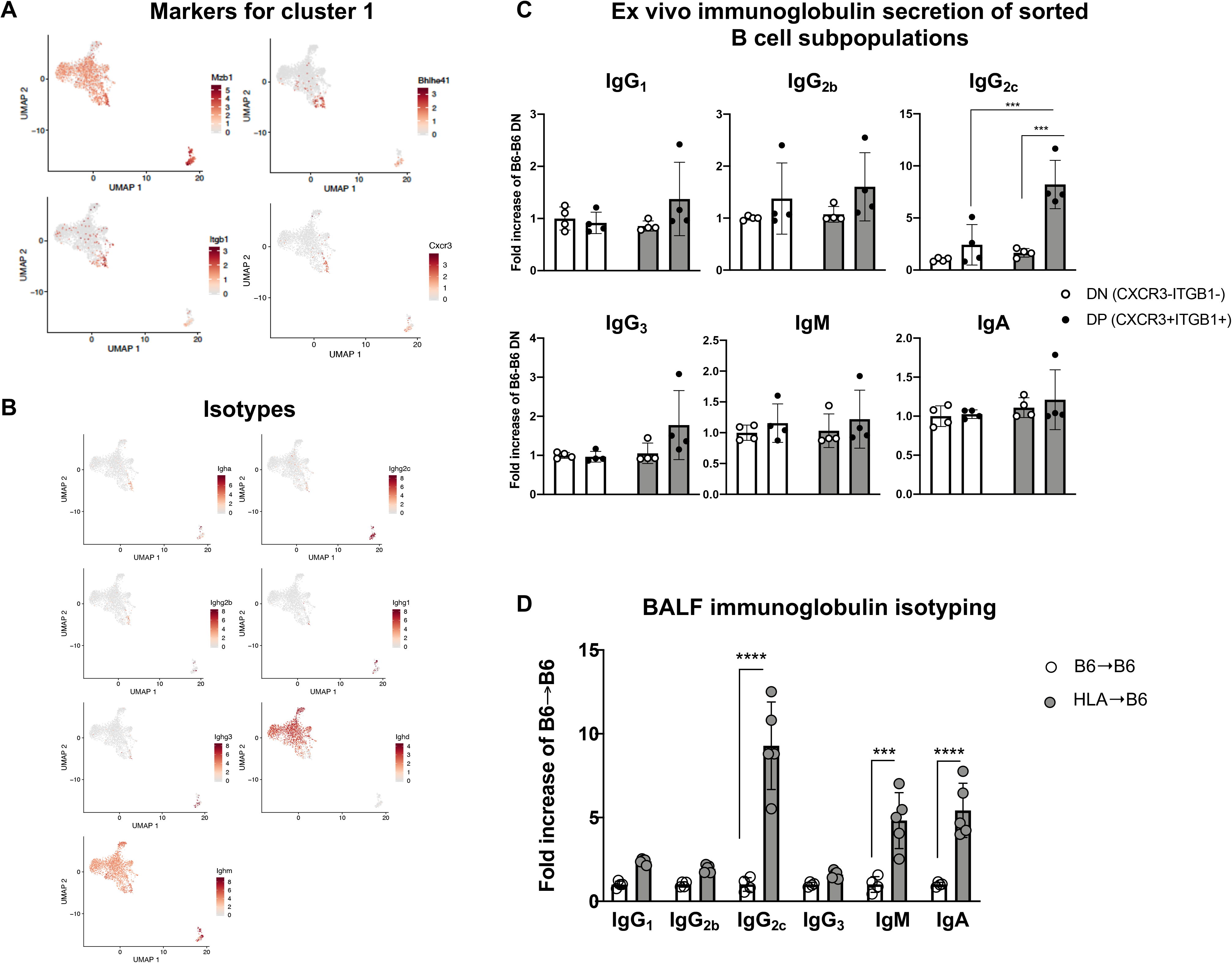
Functional characterization of the cluster #1 B cell subset. **A,** Gene expression patterns projected onto UMAP plots of *Mzb1*, *Bhlhe41*, *Itgb1*, and *Cxcr3* in the B and plasma cell subclusters from control (B6➝B6) and BOS (HLA➝B6) lung grafts. **B,** Gene expression patterns projected onto UMAP plots of immunoglobulin isotypes in the B and plasma cell subclusters from control (B6➝B6) and BOS (HLA➝B6) lung grafts. **C,** Quantification of immunoglobulin isotypes in the conditioned media from CD19+CXCR3+ITGBB1+ and CD19-CXCR3-ITGB1-cells sorted from HLA➝B6 and B6➝B6 mouse lung grafts 1 month after LTx, stimulated LPS (1 ug/mL). Data are presented as mean+/−SEM and analyzed with a 2-Way ANOVA test followed by a Tukey’s multiple comparisons test, \*\*\**p*<0.001. D, Quantification of immunoglobulin isotypes in the bronchoalveolar lavage fluid (BALF) collected from the lung grafts of HLA➝B6 and B6➝B6 mice 1 month after LTx. Data are presented as mean+/−SEM and analyzed with a 2-Way ANOVA test followed by a Tukey’s multiple comparisons test, \*\*\**p*<0.001, \*\*\*\**p*<0.0001.

### Causal role of non-IgM antibodies in the development of BOS in vivo

The observations of increased local antibody production in humans and mice with BOS prompted us to investigate their functional role, which is still subject to debate. *Aicda^−/−^* mice(24), genetically deficient for the AID recombinase and thus in the production of all immunoglobulins except IgM, received HLA grafts, and the lung function and histology of the grafts were investigated 1 month after LTx. After clamping the right bronchus, the specific lung function of the left lungs was measured. Naïve non-transplanted *Aicda^+/−^* littermates served as controls. Lung function was significantly worsened in the HLA➝*Aicda^+/+^* mice compared to naïve non-transplanted controls (**Fig. 5A**). While the compliance, elastance and IC were not different between HLA➝*Aicda^+/+^* and HLA➝*Aicda^−/−^* grafts, the resistance was lower in HLA➝*Aicda^−/−^* grafts (**Fig. 5A**). Consistently, the peribronchial fibrosis was attenuated in HLA➝*Aicda^−/−^* grafts compared to HLA➝*Aicda^+/+^* (**Fig. 5B**). The numbers of CC10+ cells however remained unchanged between HLA➝*Aicda^+/+^* and HLA➝*Aicda^−/−^* grafts (**Fig. 5C**). HLA➝*Aicda^+/+^* and HLA➝*Aicda^−/−^* grafts displayed comparable infiltrations of T and B cells around the airways (**Sup. Fig. 4**) Altogether, our results show that the prevention of the production of all the antibodies except IgM has a beneficial impact on lung graft resistance, potentially through a decrease in fibrogenesis around the airways.

**Figure 5.**
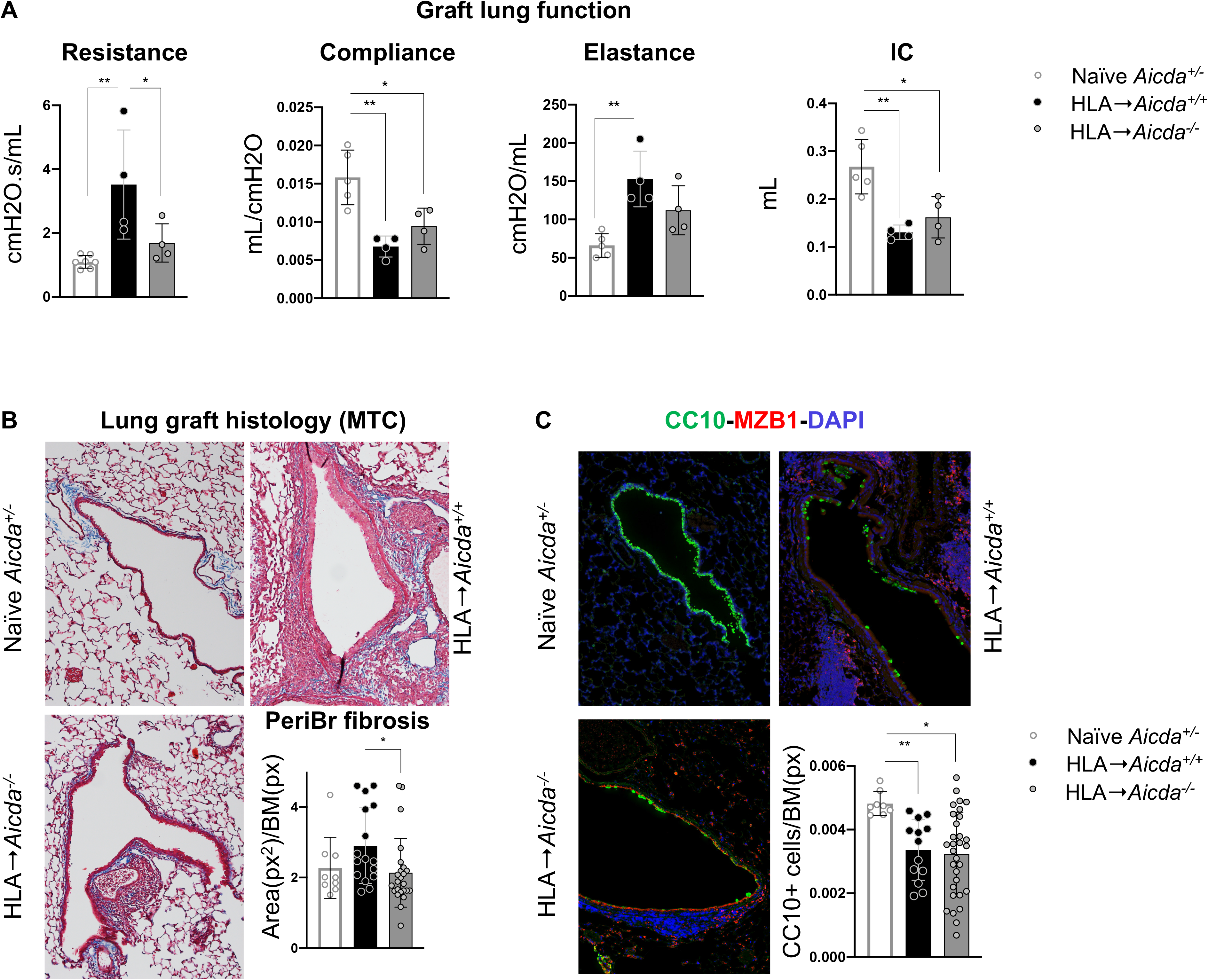
Genetic deficiency in antibody production partially protects murine lungs against the development of BOS signs. Lung grafts from HLA donors were orthotopically transplanted into *Aicda^+/+^* and *Aicda^−/−^* littermates on a B6 background (HLA➝*Aicda^+/+^* and HLA➝*Aicda^−/−^*) and analyzed 1 month later. Naïve non-transplanted *Aicda^+/−^* mice were used as baseline controls. **A,** Measurement of the lung function of the left lung grafts after clamping the right bronchus. **B,** Representative histology after Masson Trichrome (MTC staining) and quantification of the peribronchial fibrosis area, normalized to the basement membrane length. **C,** Immunofluorescence staining for MZB1+ and CC10+ cells, and quantification of CC10+ cells normalized to the basement membrane length. Data are expressed as mean+/−SEM and analyzed with a 1-Way ANOVA followed by a Tukey’s multiple comparisons test, \**p*<0.05, \*\**p*<0.01.

## Discussion

We applied high-dimensional single cell analysis with unsupervised clustering to identify the cell subsets infiltrating murine lung grafts during the progression of chronic rejection after orthotopic LTx. This analysis generated 14 large cell clusters and identified these as 11 known cell types. Subclustering of the lung structural cells resulted in 11 additional clusters. We identified cell types previously detected in the mouse lung by the scRNA-seq technique (25), and observed dramatic shifts in the relative abundance of both immune and structural cell type populations between BOS and control transplanted lungs. The gene expression signatures of these clusters offer molecular determinants in the progression of chronic rejection to BOS after LTx.

During BOS, plasma cells (PCs), identified by the expression of the classical PC markers *Xbp1*, *Sdc1*, *Mzb1*, *Irf4*(26), are the cell type exhibiting the most prominent numeric increase compared with control mouse lung grafts. Accumulations of MZB1+ PCs were consistently detected around the airways of explanted lung samples from patients with BOS and mice after LTx. While we occasionally detected PCs in lung tissue explanted from healthy donors, these were much less abundant and only positive for IgA, while PCs in BOS samples were mostly IgG+. PCs represent the terminal stage of B cell differentiation, and are known for their ability to produce high amounts of antibodies (27). We showed that total IgG levels are increased in BALF of patients with BOS after LTx, compared with patients who remained BOS-free. This finding, combined with the presence of IgG+ PCs around BOS airways, suggests a local origin of the soluble IgG measured in BALF, highlighting that a local antibody repertoire drives transplant rejection (28, 29). This significant increase of overall IgG levels in association with BOS is consistent with previously published data (30), yet extend those to increased detection of individual Ig isotypes IgG_1_, IgG_2_, IgG_3_ and IgG_4_).

While an association between antibody levels (peripheral and local) and BOS has been reported by us and others, their functional contribution to disease onset and progression remained unknown. To mechanistically address this question, we performed orthotopic LTx in *Aicda^−/−^* mice, which lack secretion of all antibody types except IgM (which was unchanged during chronic rejection). We determined that the absence of those antibody subclasses in *Aicda^−/−^* recipient mice was sufficient to prevent peribronchial fibrosis and thus alleviated the pathologic decrease of graft airway resistance after LTx. Further studies will need to determine whether the antibody functions in this context are Fc-dependent (complement-mediated cytotoxicity, FcγR-dependent inflammation) (31) or Fab-dependent (direct activation of target cells through endogenous surface proteins, which can be either MHC molecules or surface autoantigens) (32–35).

The PCs detected in lungs during BOS development were numerous around fibrotic and partially obliterated airways. We hence sought to identify the potential cellular origin of these peribronchial PCs, by subclustering B cells and PCs identified by scRNA-seq of mouse lung grafts (Fig. 3). We identified the likely cellular origin of PCs in rejected lung grafts using trajectory and pseudotime analysis. These PC progenitors were defined by a set of transcripts, encoding the cell surface markers integrin beta 1 (*Itgb1*) and Cxcr3 (known to regulate cell adhesion (36) and chemotaxis (37, 38)), as well as the transcription factors Bhlhe41, Zbtb20, and Zbtb32. In murine lung grafts, both the PC cluster, as well as their putative progenitors, were the only intragraft B cells positive for genes encoding IgA, IgG_1_, IgG_2b_, IgG_2c_ and IgG_3_. In BALF, IgA and IgG_2c_ were significantly increased compared with controls. IgA, mostly produced around the bronchial epithelium in steady state, is considered protective, as it is the first line of defense against airborne pathogens (39). IgA is therefore unlikely to participate to rejection after LTx, although in some rare cases, IgA has been found capable of contributing to autoimmunity, as in the case of IgA nephropathy (40). We sorted CXCR3+ITGB1+ PC progenitors from mouse lung grafts and demonstrated that this population specifically contributed to the increased production of IgG_2c_. The IgG_2c_ is mainly generated in response to carbohydrates and proteins (31) and provides protection against viruses (41, 42), but its contribution to chronic rejection was unknown up-to-now.

Using the ImmGen database (23), we determined that PC progenitors identified in rejected mouse lung grafts display a B-1 phenotype. This is based on their expression of *Bhlhe41*, a transcription factor recently identified as a marker of B-1 cells, shaping their development, homeostasis, and BCR repertoire (22), as well as the classical B-1 markers Cd43 and IgM. In addition to these conventional markers, this BOS-associated B-1 cell population displays markers previously undefined or unknown in the characterization and/or function of B-1 cells, such as Cxcr3 and Itgb1. The transcriptome of the chronic rejection-associated B-1 cell population did not display a major overlap with innate response activator B cells, another B-1 cell population, that was discovered to regulate several pathological contexts (**Sup. Fig. 5**)(43).

Our data hence identify a novel role of the B-1 cell population in BOS (44–46). B cells are classically subdivided in conventional B-2 cells and innate-like B-1 cells (47, 48). Conventional B-2 cells are the prototype of adaptive humoral immunity, selected for the production of antibodies with the highest specificity and affinity against encountered antigens. While the involvement of B-2 cells in infectious and chronic inflammatory diseases has been extensively investigated, the role of B-1 cells in disease has been underappreciated. Innate B-1 cells are distinguished from B-2 cells by their different developmental origin, a self-renewal capacity, *in situ* location, and a number of phenotypical and functional characteristics. While the antibody repertoire of B-2 cells constantly evolves upon antigen encounters, the B-1 repertoire in mostly predetermined during prenatal development (49). B-1 cells produce polyreactive antibodies (low affinity and broad specificity) targeting common pathogenic motifs or intracellular autoantigens (49). B-1 cells secrete natural IgM, but recent evidence, including our findings, show that they can also switch antibody classes and produce IgG for instance (50). While B-1 cells and their natural IgM show beneficial effects in e.g. atherosclerosis (51), the intrinsically autoreactive nature of their antibody repertoire also assigns to them a recently described role in autoimmunity (52), in particular due to the generation of autoantibodies in transplantation (52, 53). Autoimmunity is associated with BOS after LTx, as evidenced by increased production of anti-Collagen V autoantibodies and their potential to induce BOS (54, 55). Our data suggest that a subset of innate B-1 cells, identified by classical and context-specific markers (Bhlhe41, Cd43, IgM and Cxcr3, Itgb1, respectively), can contribute to autoimmunity in BOS, and thus represent potential therapeutic value. Polyreactive antibodies have recently been associated with cardiac allograft vasculopathy after heart transplantation (56, 57). More recently, innate B cells have been detected in rejected human kidney grafts (58). To the best of our knowledge, this is the first report that identifies a set of unique markers for this innate B cell population and unravels a mechanistic contribution of this local innate B cell subpopulation to chronic graft dysfunction and rejection. Importantly, in addition to the production of potentially pathogenic antibodies, these subpopulations of B-1 cells can also produce protective IgM (59), regulate inflammatory processes (46) and T cell differentiation and function (48), or even directly interact with structural cells to impacting their function in disease (60). In conclusion, our results identify a Cxcr3+Itgb1+ B-1 cell subset in lung grafts undergoing chronic rejection after LTx, which serve as progenitors to intra-graft PCs capable of producing high levels of IgG_2c_ in association with the development of BOS.

## Materials and methods

### Patient demographics

USF-TGH LTx cohort. Samples of bronchoalveolar lavage fluid from patients who developed or not BOS after LTx were from a sample of the Tampa General Hospital LTx cohort (IRB# Pro00032158, University of South Florida, USA). The demographics are presented on Table 1.

University of Colorado collection. Paraffin-embedded samples of human BOS lung tissue were from the Department of Pathology Department at the University of Colorado – Anschutz Medical Campus. The tissue was explanted from 4F and 6M patients who underwent a lung transplantation and received a second transplant at the UCH (CO, USA) after they developed end stage BOS (IRB# 18-0178, Colorado Multiple Institutional Review Board, COMIRB). Paraffin-embedded samples of healthy lung tissue were collected from dead donors at the National Jewish Hospital, CO, USA (IRB# 11-1664, COMIRB).

### Orthotopic lung transplantation in mice

Male C57BL/6 (B6), and HLA (*C57BL/6-Tg(HLA-A2.1)^1Enge/J^*) mice were purchased from The Jackson Laboratory. *Aicda^−/−^* mice were a gift from Dr. Jing Wang from the University of Colorado – Anschutz Medical Campus. Lung transplantations (LTx) were performed as described previously(4) with minor modifications. No immunosuppression was applied to any of the transplanted mice. B6 and HLA mice were used as donors, and B6 as recipients. Briefly, donors were anesthetized with an i.p. injection of ketamine/xylazine. The pulmonary artery, bronchus, and pulmonary vein were carefully separated from one another with blunted forceps, prior to cuffing with, respectively, 24−, 20−, and 22-gauge cuffs. The left lung graft was stored for less than 1 hour before its implantation. The recipient mouse was anesthetized with a mixture of medetomidine (1 mg/kg), midazolam (0.05 mg/kg), and fentanyl (0.02 mg/kg); intubated; and connected to a small-animal ventilator (Harvard Apparatus), at a respiratory rate of 120 bpm and a tidal volume of 300 μl. The chest was opened on the left side between ribs 3 and 4, and the native left lung was retracted with a clamp. The hilar structures were carefully separated from one another with blunted forceps. After arrest of the blood and air flow toward the left lung, the cuffed graft pulmonary artery, bronchus, and pulmonary vein were inserted into the recipient counterparts and ligated with 9-0 sutures. The native left lung was removed, and the incision in the chest was closed with a 6-0 suture, after removing all potential air bubbles from the chest. Antagonist was administrated, and the animal was extubated when it showed signs of spontaneous breathing. After the operation, the recipient mice were allowed to recover at 30°C overnight and received a single injection of buprenorphine SR (1 mg/kg). The mice were sacrificed at one-month post-LTx. All animal experimentations were approved by the IACUC of the University of Colorado (protocol 115517(04)2D).

### Preparation of murine lung grafts as single cell suspensions

Single cell suspensions of murine lung grafts were used for the single cell RNA-sequencing analysis, flow cytometry and fluorescence-activated cell sorting. The graft lung tissue was digested and separated into single-cell suspensions using the Lung Dissociation Kit, Mouse (Miltenyi) according to the manufacturer’s instructions.

#### Single cell RNA-sequencing

Single cells were captured using the 10x genomics chromium platform. 10,000 cells were loaded and single cell libraries were generated using the 10x genomics 3’ end gene expression kit.

#### Single cell RNA-seq analysis

Sequencing data was processed using the cell ranger pipeline from 10x Genomics with the mm10 genome assembly to generate unique molecular identifier (UMI) count matrices for each sample. Seurat was used to perform normalization, clustering, and to generate UMAP projections(61). The R package SoupX was used to remove free RNA contamination(62). Principal component analysis was performed on z-score transformed data using 30 dimensions for the full dataset and 20 dimensions for the structural lung cells and B-cell subsets. Graph-based clustering was performed using a resolution of 0.1, 0.3, and 0.1 for the full dataset, structural cells, and B-cell subsets respectively. The R package clustifyr was used to aid in assigning cell types to clusters (63). The Tabula Muris mouse single cell RNA-seq dataset was used to define a reference dataset for calling cell types (11).

Genes differentially expressed in each cluster compared with other clusters in each tested comparison were determined with a Wilcox rank sum test from the Presto R package (64). Heatmaps were generated using ComplexHeatmap or Seurat (65). Pseudotime values were calculated using the CytoTrace algorithm and RNA velocity vectors were computed and plotted using the velocyto Python package (66).

#### Data/code availability

Sequencing data, UMI count matrices, and cell level meta-data will be deposited in the NCBI’s Gene Expression Omnibus database (https://www.ncbi.nlm.nih.gov/geo/). Analysis scripts and an interactive UCSC cell browser instance will be provided at a GitHub repository (https://github.com/).

### Measurement of immunoglobulins in human bronchoalveolar lavage fluid

Immunoglobulins from human BALF were measured with the LEGENDplexTM Human Immunoglobulin Isotyping panel (BioLegend), according to the manufacturer’s instructions.

### Flow cytometry

Single cell suspensions from murine lung grafts were counted, then incubated with the following antibodies: anti-mouse CD19-PerCP Cy5.5 (eBioscience, Thermo Fisher), anti-CD183(CXCR3)-PE, mouse (Miltenyi Biotec), anti-CD29(ITGB1)-APC, mouse (Miltenyi Biotec), PE anti-mouse CD43 (BD Pharmingen), anti-mouse CD5-APC (eBioscience, Thermo Fisher). Fluorescence was quantified using a FACS Canto II flow cytometer (BD Biosciences) and analyzed with the DIVA software (BD Biosciences).

### Fluorescence-activated cell sorting and ex vivo stimulation of B cell subpopulations

Single cell suspensions from murine lung grafts were counted, then incubated with the following antibodies: anti-mouse CD19-PerCP Cy5.5 (eBioscience, Thermo Fisher), anti-CD183(CXCR3)-PE, mouse (Miltenyi Biotec), anti-CD29(ITGB1)-APC, mouse (Miltenyi Biotec). The cells were sorted into cooled sterile PBS+10%FBS, into two cell populations: CD19+CXCR3+ITGB1+ and CD19+CXCR3-ITGB1-using the BD Aria I cell sorter (BD Biosciences). The cells were centrifuged at 300g, then resuspended in RPMI, supplemented with 10% FBS, penicillin/streptomycin, 2 mM glutamine, 0.1 mM nonessential amino acids, 1mM sodium pyruvate and 50 uM 2-mercaptoethanol. The cells seeded in a round-bottom 96 well-plate, and cultured for 5 days with LPS (1 ug/mL) at 37C in 5% CO2. For the measurement of immunoglobulins, the supernatants were collected.

### Measurement of murine immunoglobulins

Immunoglobulins were measured in preconditioned media from LPS-stimulated B cell subpopulations, sorted from murine lung grafts. The levels of IgG2c were measured using the IgG2c Mouse ELISA kit (Thermo Fisher), according to the manufacturer’s instructions. The levels of IgG1, IgG2b, IgG3, IgA, IgM and IgD were measured using the Mouse Immunoglobulin Isotyping kit (BD Biosciences), according to the manufacturer’s instructions.

### Histology and immunostaining

Sections of paraffin-embedded specimens were acquired from the Pathology core of the University of Colorado. Mouse lung grafts were fixed with PFA 4% instilled intratracheally for 24h. The specimens were embedded in paraffin and prepared as 4um-thick sections at the Histology Core of the National Jewish Hospital. Both human and mouse paraffin sections were deparaffinized by melting the paraffin at 60C, followed by incubations in sequential xylene, ethanol and water baths. Antigen retrieval was performed in 0.01 M citric acid, pH 6.0, in a pressure cooker for 30 seconds at 125°C. For immunohistochemistry staining of MZB1, endogenous peroxidases were inhibited with 3%H2O2, then the Avidin/Biotin blocking kit (Vector laboratories) was used. Sections were blocked with PBS+3%BSA, then incubated with the primary anti-MZB1 antibody (polyclonal, Sigma-Aldrich) overnight at 4C. Sections were incubated with a secondary biotinylated anti-rabbit antibody (Agilent Technologies), followed by peroxidase (ABC Vectastain kit, Vector laboratories). MZB1 was finally detected with the DAB substrate (Vector laboratories). The image acquisition was performed with the Aperio slide scanner (Leica Biosystems), and the eSlide manager software. For immunofluorescence staining, sections were blocked with PBS+3%BSA, then incubated overnight at 4C with the following primary antibodies: anti-MZB1 (Sigma-Aldrich), anti-CD38 (Santa-Cruz Biotechnology), anti-IgG (clone EPR4421, Abcam), anti-IgA (clone EPR-5367-76, Abcam). Finally, the sections were incubated with fluorochrome-conjugated secondary antibodies (Invitrogen). The images were acquired with an Olympus microscope.

### Left lung graft lung function measurements

Mice were anesthetized with pentobarbital, then intubated and connected to a Minivent-ventilation system (120 breaths/min, volume of 300μL). The chest was opened, and the right bronchus was clamped. Then, the left lung function was measured with a FlexiVent system (Scireq) (tidal volume of 3.75 mL/kg, frequency of 150 breaths/min). Airway compliance, resistance, tissue elastance and inspiratory capacity were measured using the SnapShot, Prime-8 and Quick-prime wave perturbations. Three readings per animal were taken.

### Statistical analysis

Data represent mean ± SEM, from n separate experiments. Statistical significance of differences was evaluated by a Mann-Whitney unpaired 2-tailed t test or a 1-way ANOVA followed by a Tukey’s multiple comparisons post-test or a 2-way ANOVA followed by a Tukey’s multiple comparisons post-test. Differences were considered to be statistically significant at *p*<0.05. The types of analysis used for each data set are indicated in the legends of the corresponding figures.

## Acknowledgements

We would like to acknowledge the Immunological Genome Project (ImmGen).

