## Supplementary Material for "Single cell transcriptome mapping identifies a local innate B cell population driving local antibody production and chronic rejection after lung transplantation"

**Supplementary Figure Legends**

**Sup. Fig. 1. Annotation of cell populations from mouse lung grafts. A,** Number of cells included in the single cell RNA-sequencing analysis per population and per sample. **B,** Heatmap for cell annotations of the different clusters after correlation to the gene expression profiles of the Tabula Muris (1). **C,** Uniform Manifold Approximation and Projection (UMAP) representation of 11 distinct cell populations from unsupervised clustering of gene expression data in single cells extracted from control (B6→B6) and BOS (HLA→B6) transplanted mouse lungs. Selected marker gene expression is represented according to the provided color scheme. **D,** Violin plots of log-transformed gene expression of selected cell population marker genes displayed for the control (B6→B6) and BOS (HLA→B6) transplanted mouse lung conditions.

**Sup.Fig. 2. Analysis of structural lung cells from mouse lung grafts.** The cell population identified as structural lung cells in the analysis of control (B6→B6) and BOS (HLA→B6) lung grafts was submitted to unsupervised subclustering. **A,** Uniform Manifold Approximation and Projection (UMAP) representation of 11 distinct cell populations from unsupervised clustering of gene expression data in single cells identified as structural lung cells extracted from control (B6→B6) and BOS (HLA→B6) transplanted mouse lungs (top). UMAP representation of the cell populations detected in murine lung grafts by condition (HLA→B6 and B6→B6) (bottom left). Proportion of each cell type in control vs. BOS mouse lung graft samples (bottom right).

**Sup. Fig. 3. Flow cytometry analysis of lung graft B cell populations.** 1 month after lung transplantation (LTx), mouse lung grafts from control (B6→B6) and BOS (HLA→B6) were dissociated into single cells and the CD19+CXCR3-ITGB1- (non-cluster #1) and CD19+CXCR3+TGB1+ B cells were analyzed and sorted by FACS (top). The expression of CXCR3, ITGB1 and classical B-1 cell markers CD43, CD5 was confirmed after a 5 day ex vivo culture of the sorted B cell populations (bottom).

**Sup. Fig. 4. T cell and B cell infiltrations around the bronchi are unchanged in the absence of Aicda expression in lung transplant recipient mice.** Lung grafts from HLA donors were orthotopically transplanted into *Aicda*$^{+/+}$ and *Aicda*$^{-/-}$ littermates on a B6 background (HLA→*Aicda*$^{+/+}$ and HLA→*Aicda*$^{-/-}$) and analyzed 1 month later. Representative pictures of bronchial areas after immunofluorescence staining for CD45R+ B cells and CD3+ T cells.

**Sup. Fig. 5. Comparison of lung graft B cell subsets to available data.** Data from the single cell RNA-sequencing analysis of B and plasma cell cluster identified in the mouse lung grafts was compared to the B cell subsets identified by Rauch and colleagues and deposited in the NCBI GEO database (series GSE32372), and presented as a correlation.

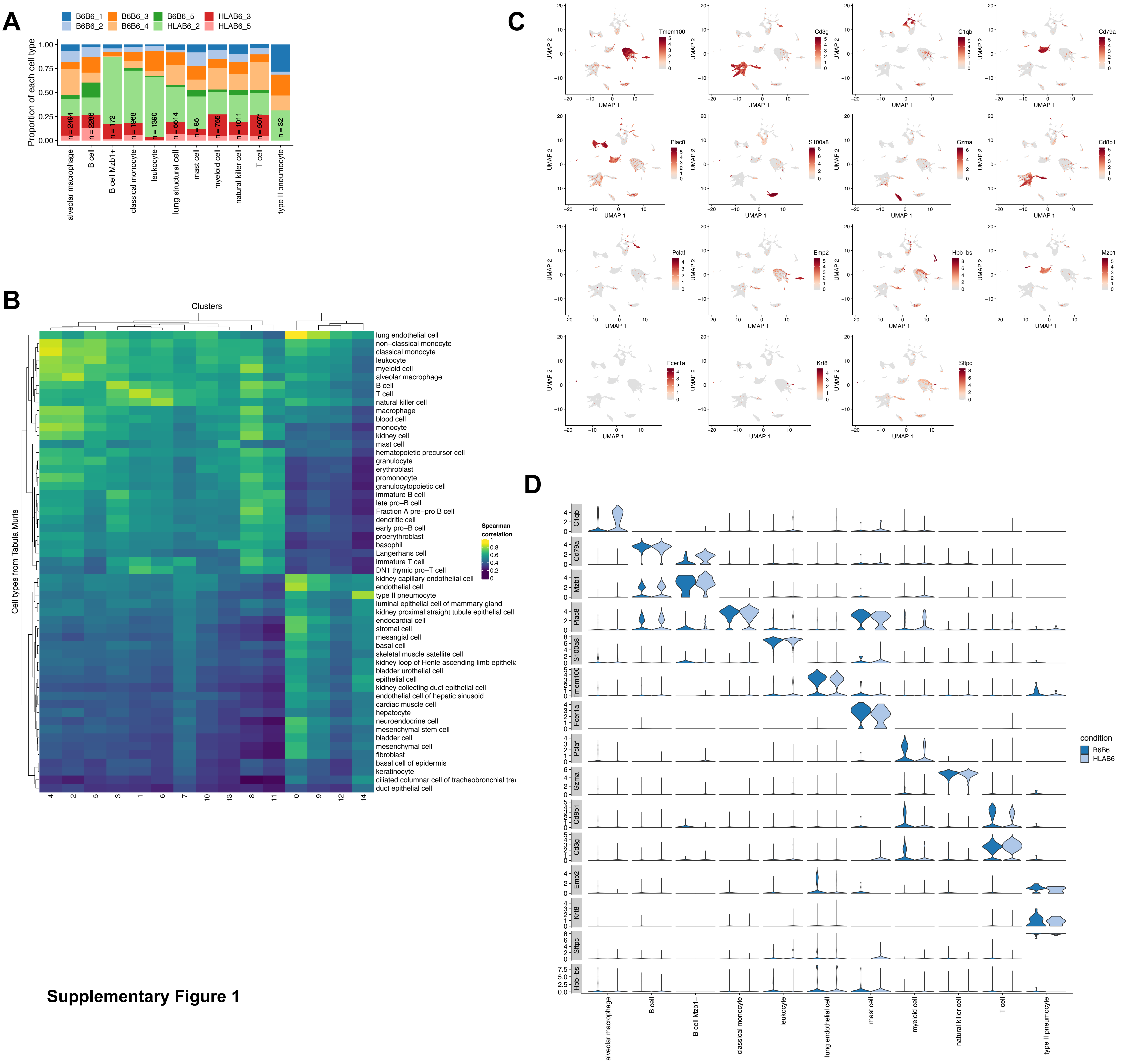


**Supplementary Figure 1**

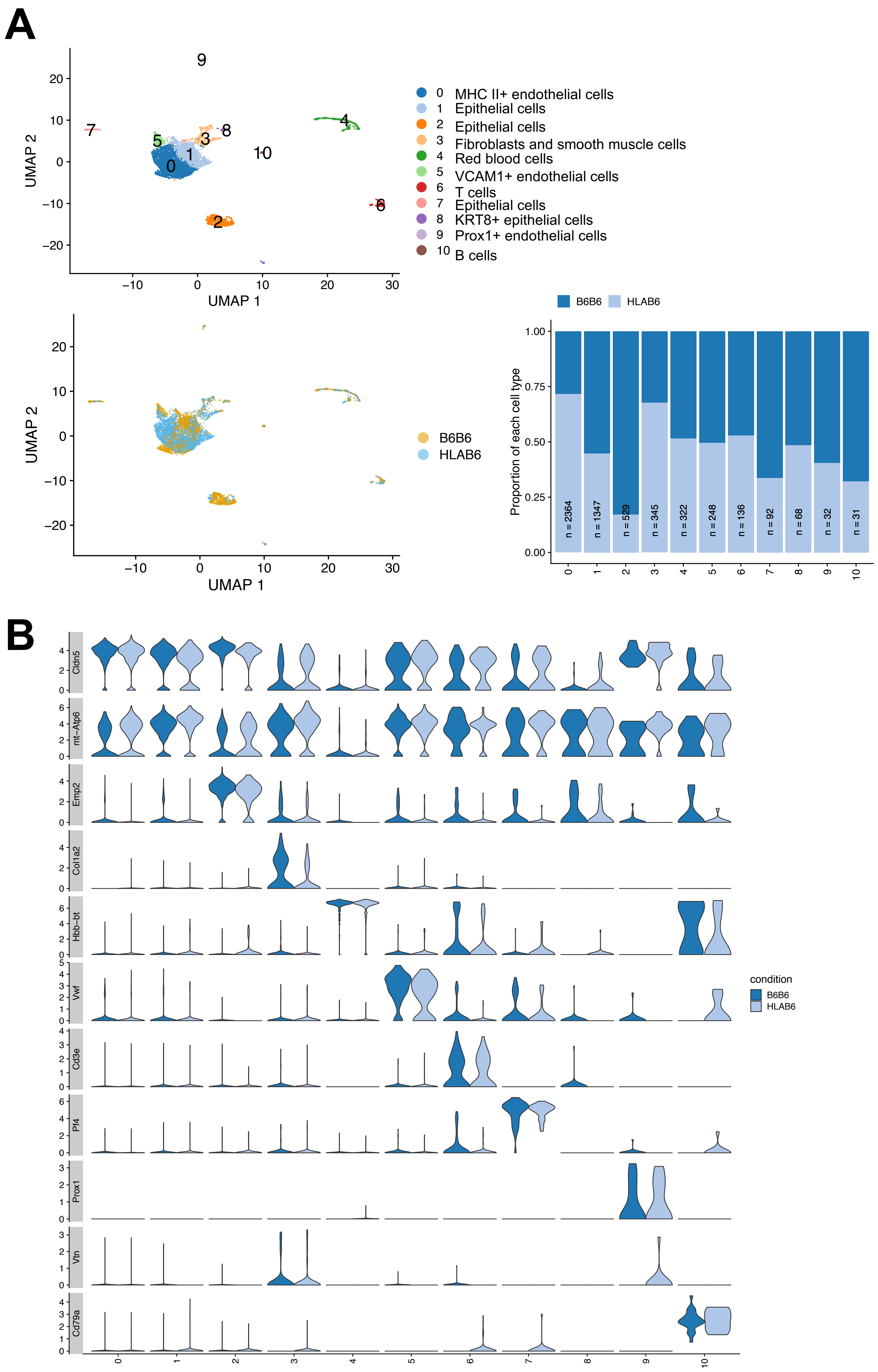


### Supplementary Figure 2

### Flow cytometry analysis of lung graft B cell populations

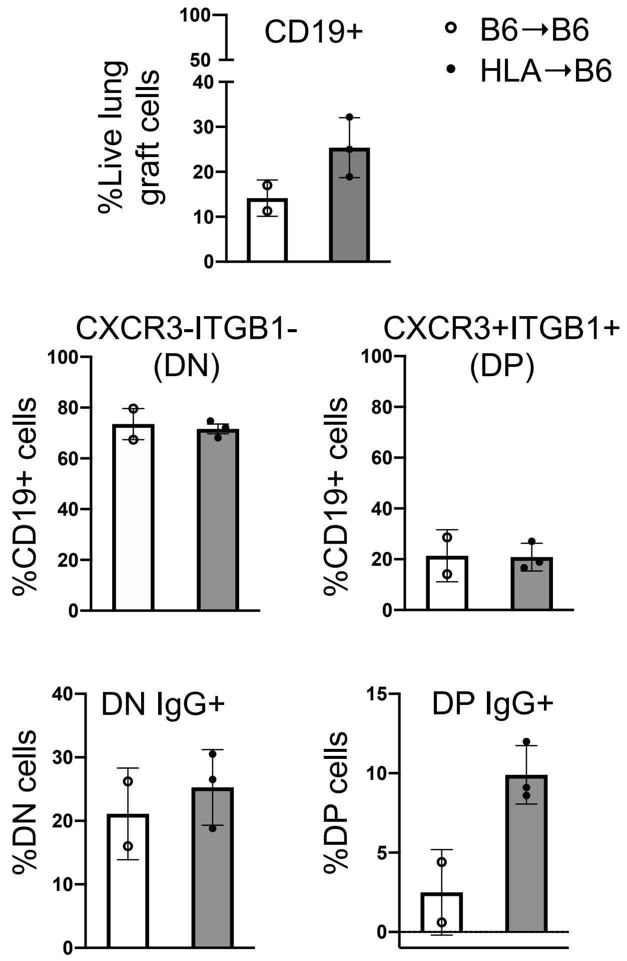


### Sorted B cell subpopulations

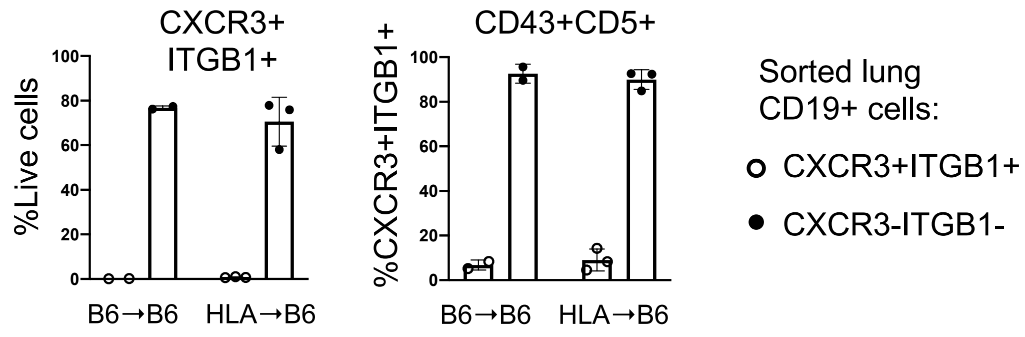


**Supplementary Figure 3**

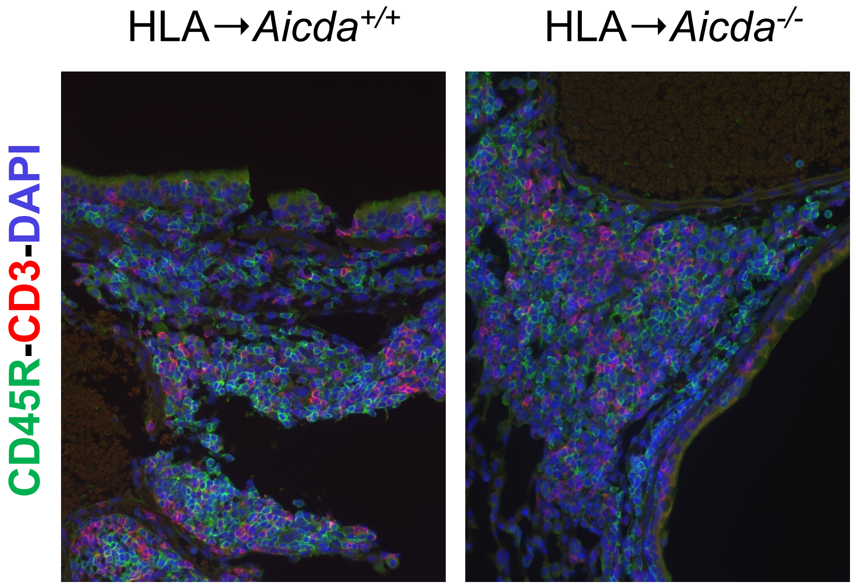


**Supplementary Figure 4**

**Comparison of murine lung B cell clusters to B cell subsets of GSE32372 GEO database**

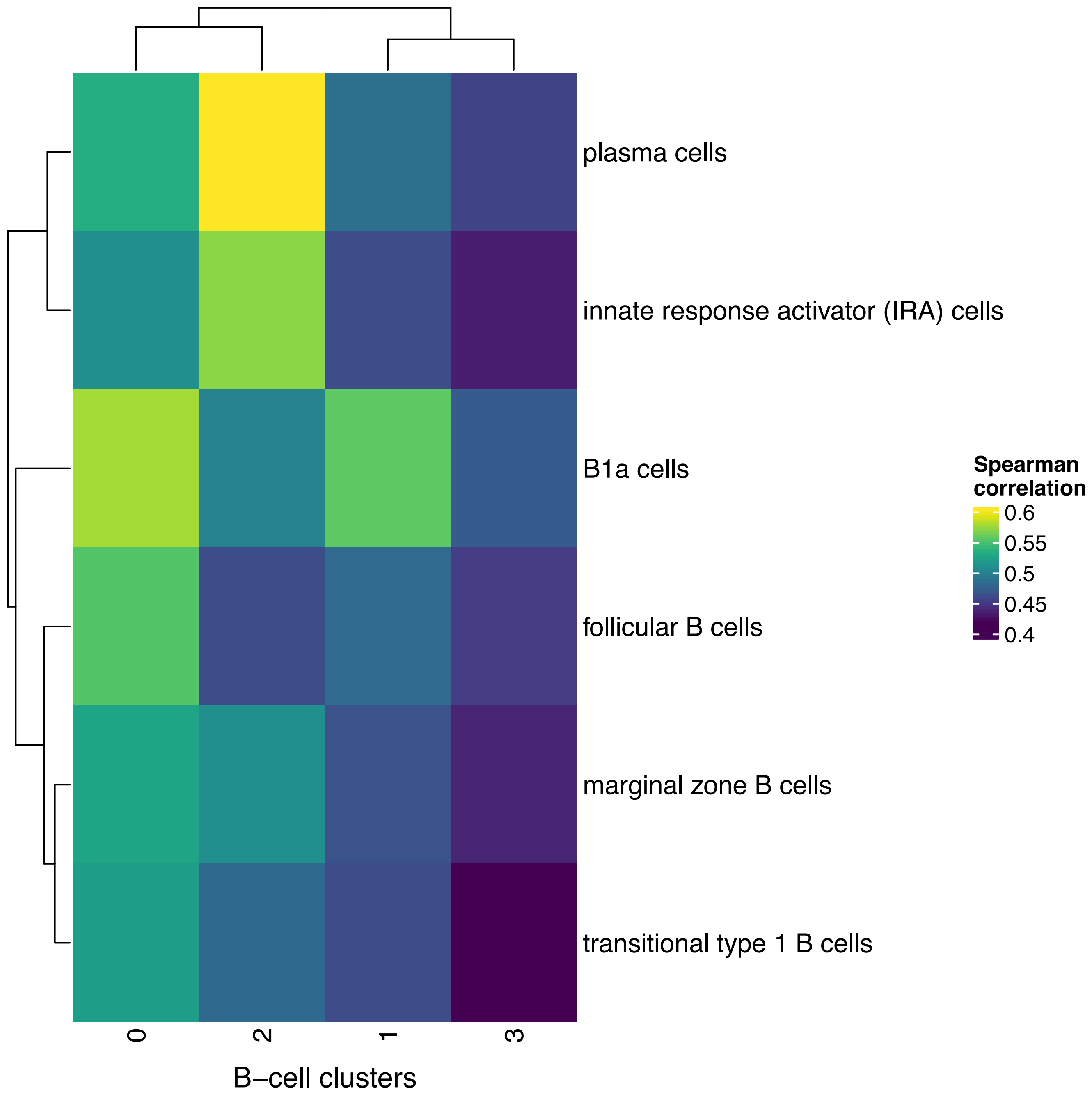


**Supplementary Figure 5**
